# RNA modulates FUS condensate assembly, dynamics, and aggregation through diverse molecular contacts

**DOI:** 10.64898/2025.12.13.694118

**Authors:** Tongyin Zheng, Kandarp A. Sojitra, Samara Cummings, Qizan Chen, Priyesh Mohanty, Jeetain Mittal, Nicolas L. Fawzi

**Author notes:** Correspondence should be addressed to: J.M. and N.L.F. T.Z. and K.A.S. contributed equally to this work.

## Abstract

Fused in sarcoma (FUS) is an RNA-binding protein that undergoes phase separation with RNA and other cellular components, forming ribonucleoprotein (RNP) granules. While recent advances delineating the molecular forces that underlie phase separation have largely focused on protein–protein interactions (1–6), the molecular details of protein-RNA interactions within condensates remain limited. In this study, we demonstrate that RNA modulates the phase separation of the low-complexity (LC) and arginine-glycine-glycine motif (RGG1) domains of FUS: low RNA concentrations enhance protein phase separation and excess RNA disrupts it. By integrating biochemical assays, NMR spectroscopy, and molecular dynamics simulations, we show that RNA incorporates into FUS condensates, reducing condensate density while enhancing local relaxation and diffusional motion of FUS. Surprisingly, whereas RNA binding in the dispersed phase primarily involves the RGG1 domain, within the condensed phase, both LC and RGG1 domains contribute to interactions with RNA. NMR and simulation data show diverse interactions between amino acids and RNA moieties, including prominent glutamine-RNA contacts, that stabilize FUS-RNA co-condensates. Furthermore, we found that RNA accelerates the liquid-to-solid transition of FUS LC-RGG1 condensates, promoting fibrillar aggregate formation. Together, these results provide mechanistic insight into how RNA regulates the assembly, dynamics, and maturation of protein condensates.

Graphical Abstract
Using NMR and molecular simulations, we map how RNA engages FUS LC-RGG1 within condensates through electrostatic, π-stacking, and hydrogen-bond contacts. We find that RNA incorporation dilutes condensate density, tunes protein mobility, remodels interaction networks, and accelerates the formation of fibrillar aggregates.

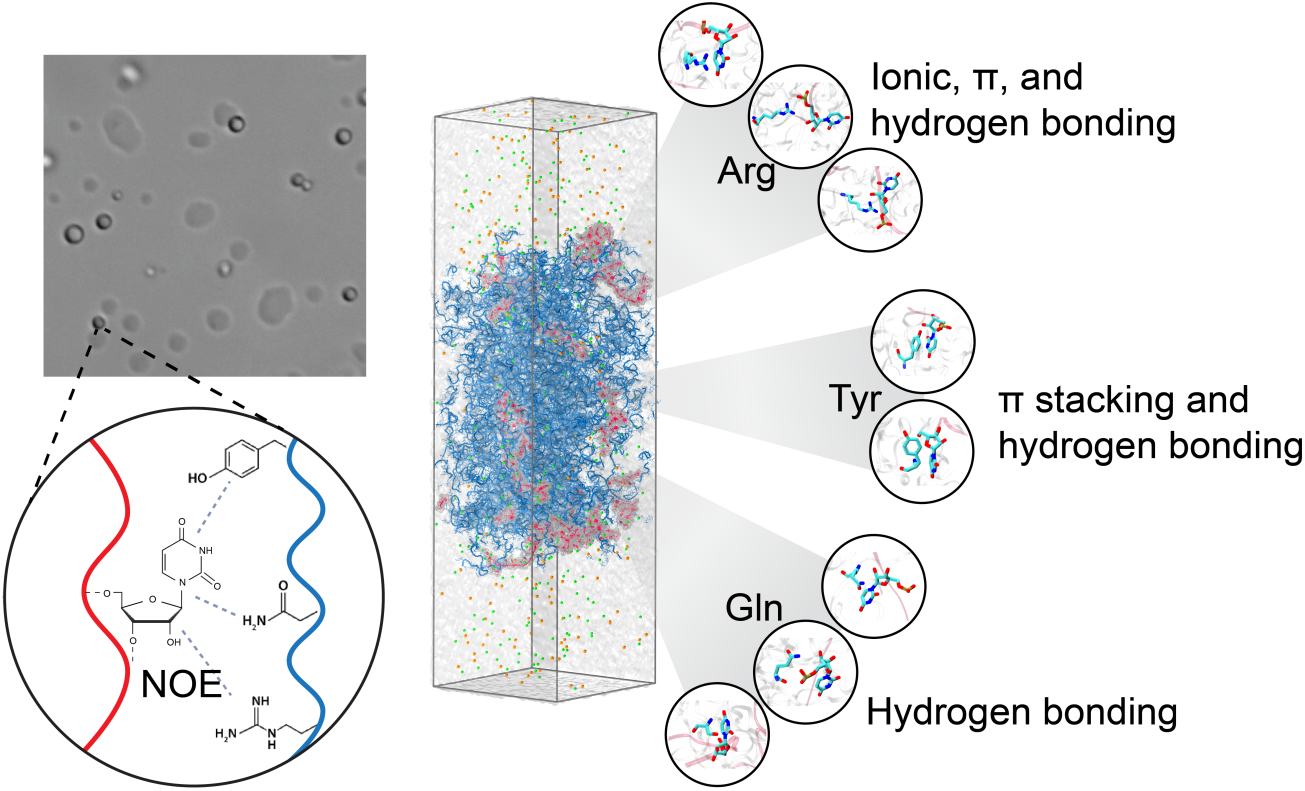

## Introduction

Ribonucleoprotein (RNP) granules are specialized cellular compartments enriched with RNA and RNA-binding proteins that serve as hubs for RNA-related processes such as storage, transportation, and translation regulation (7). These non-membrane bound condensates can form through reversible phase separation of biomolecules, allowing them to assemble and disassemble rapidly in response to cellular signal or environmental changes (8). Importantly, aberrant transition of RNP granules and their constituent proteins are widely believed to be connected to the pathogenesis of neurodegenerative diseases (9).

Fused in Sarcoma (FUS) is an RNA-binding protein found in stress granules (SGs), a cytoplasmic RNP granule that forms in cellular stress response. Dysregulation of FUS, manifested as FUS-positive cytoplasmic aggregates, has been linked to neurodegenerative diseases, particularly amyotrophic lateral sclerosis (ALS) and frontotemporal lobar degeneration (FTLD) (10–12). FUS harbors multiple RNA-interacting domains and regions, including two folded domains: an RNA recognition motif (RRM) and a zinc finger (ZnF), and three disordered arginine-glycine-glycine motif (RGG) domains. The RRM and ZnF bind RNA with specific sequence and structural preferences, while the RGG domains are believed to mainly mediate non-specific interactions with RNA and enhance the binding affinity of the RNA recognition motifs (RRMs) (13). Additionally, FUS contains an N-terminal low complexity domain (LC), which is enriched in polar residues and responsible for driving the phase separation of FUS. Importantly, although FUS can phase separate in the absence of RNA, RNA has been shown to tune the phase separation of FUS depending on factors such as RNA length, sequence, and concentration (13–15).

To date, most existing insights into RNA interactions with proteins associated with RNP granules have been derived from the analysis of folded domain-RNA complex structures in the dilute phase. Most notably, crystallographic and/or solution NMR structural ensembles of RRMs complexed with RNA have been determined for RNPs such as TDP-43, hnRNPA1 and FUS (16–20). In complexes with folded proteins (21), RNA often engages in hydrophobic interactions involving its bases and aliphatic/aromatic residues (22–25), as well as hydrogen bonding and electrostatic interactions that can potentially involving RNA bases, ribose rings, and phosphate groups (26–28). π-interactions form between nitrogenous base ring and aromatic amino acids (Trp, Tyr, Phe, and His) as well with residues with sp^2^ hybridized side chains (Arg, Glu and Asp) and have been found with high frequency (29–31). In contrast, interactions between RNA and disordered, Arg/Lys-rich low-complexity regions, such as FUS RGGs, are generally thought to be dominated by electrostatic attraction between basic residues and the RNA backbone (21).

Recent studies have significantly advanced our understanding of protein-protein interactions within biomolecular condensates. Based on biochemical and atomically detailed biophysical studies, non-covalent, multivalent interactions between protein molecules using many interaction modes (e.g. charge-charge, hydrophobic, π-π, and hydrogen bonding interactions) have been found to underlie protein phase separation (1–6). However, our knowledge of proteins-RNA interactions inside condensates remain largely unknown, even though such interactions contribute significantly to phase separation and RNP granule formation (32, 33). Sequencing-based studies nevertheless suggest that the protein-RNA interaction landscape is altered in condensates: although some degree of binding specificity is maintained (34), the RNA interactome of RBPs like FUS differs substantially between dispersed and phase-separated states (35).

To understand the molecular basis of RNA interaction with the disordered domains of RNA-binding proteins within condensates, we employed an integrative strategy that combines biochemical experiments, solution-state NMR spectroscopy, and computational modeling. We addressed several outstanding questions: (i) how does RNA modulate the phase behavior of FUS LC-RGG1?, (ii) how do the LC/RGG1 domains (and their amino acid types) interact with RNA in the condensed phase, and how do these interactions differ from those occurring in the dilute phase?, (iii) how does RNA influence the intra-condensate dynamics of FUS LC-RGG1?, and (iv) how does RNA alter the transition of FUS LC-RGG1 from liquid-like to solid-like phases?

## Results and discussion

### FUS LC-RGG1 phase separation is modulated by RNA

Previous studies have shown that RNA can modulate the phase behavior of various RNA-binding proteins including TDP-43, hnRNPA1, and FUS (36–39). In order to study RNA/protein phase separation, we sought to establish a model system where RNA plays an integral role in mediating contacts contributing to phase separation. Here, we tested the ability of polyU RNA to induce formation and dissolution of phase separation of a long, disordered segment of FUS encompassing the N-terminal low complexity (LC) and RGG motif domains (FUS LC-RGG1, residues 1-284) (Fig. 1A). FUS LC-RGG1 is an excellent model to study the phase separation of FUS-family proteins due to its similar sequence features (composition and organization) while exhibiting a saturation concentration (*C*_sat_) of ∼10 μM, comparable to that of full-length FUS (∼1 μM) (1). Below the *C*_sat_ (5 μM), we observed by differential interference contrast (DIC) microscopy that polyU RNA induces reentrant phase separation of FUS LC-RGG1 (Fig. 1B), showing no droplets without RNA, droplets with addition of small amounts of RNA, and no droplets with addition of large amounts of RNA. To assess the extent of phase separation across RNA concentrations, we measured the turbidity of FUS LC-RGG1/RNA mixtures containing polyU RNA (Fig. 1C). Complementary to DIC microscopy, we find that RNA promotes FUS LC-RGG1 phase separation initially but at high RNA concentrations phase separation is decreased and ultimately abolished, consistent with a reentrant phase transition seen previously for other proteins (40–42). These results establish mixtures of FUS LC-RGG1 and polyU RNA as an effective model for reentrant phase separation.

**Figure 1.**
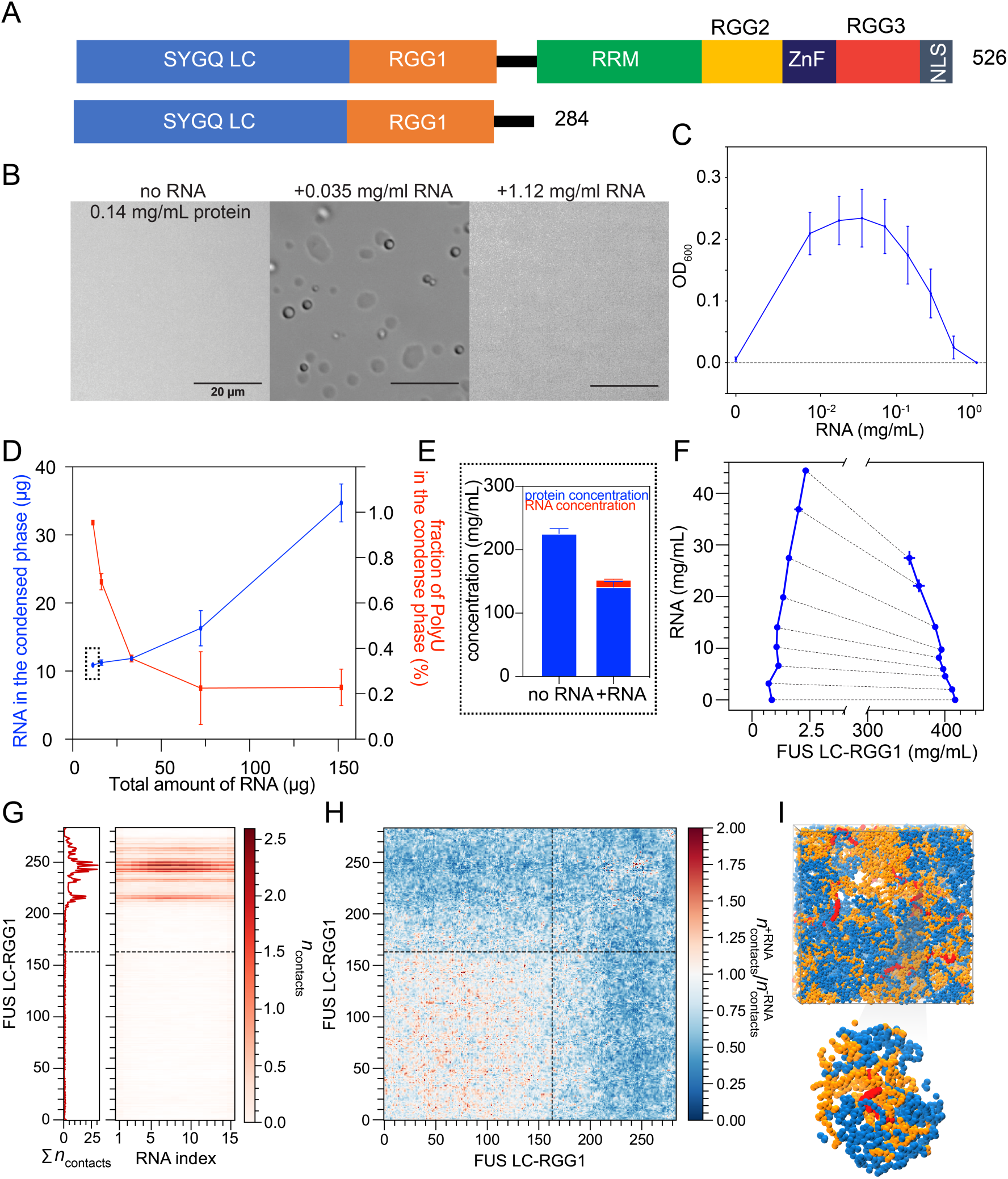
RNA modulates FUS LC-RGG1 phase separation. A) Domain architecture of FUS. FUS LC-RGG1 consists of the SYGQ LC domain and the first RGG region. B) DIC microscopy of FUS LC-RGG1 in the absence and in presence of polyU RNA of different concentrations. At low protein concentration (5 μM, 0.14 mg/ml), FUS LC-RGG1 does not phase separate (*left*) but with addition of small amounts of polyU RNA (*center*) phase separation is enhanced. Addition of larger amounts of RNA (*right*) disrupts phase separation, consistent with re-entrant phase separation modulated by RNA. C) Turbidity of FUS-RNA samples with 0.14 mg/mL protein and 0-1.14 mg/mL polyU RNA measured at 600 nm wavelength. D) Measurements of the amount of RNA incorporated into the condensed phase. The amount depends on the total RNA added to the mixture (80 μM FUS LC-RGG1, equal to 136 μg in 60 μl, with addition of 11 to 151 μg polyU RNA). As the total RNA increased, the amount of RNA incorporated also increased. However, the percentage of total RNA incorporated decreases. E) Protein and RNA concentrations of condensed phases for the first condition in D) and for a no RNA control. Incorporation of a small RNA mass fraction markedly lowers the condensed-phase protein concentration. F) Phase diagram of FUS LC-RGG1 and U_15_ RNA from CG simulation. The tie line becomes negative with the addition of RNA, indicating a reduced ability of the protein to recruit RNA. G) Per chain intermolecular contacts between FUS LC-RGG1 and RNA from bulk dense phase CG simulations. Bottom panel shows the average number of contacts formed per residue. Vertical dashed line indicates the LC-RGG1 domain boundary. H) Per chain intermolecular protein-protein contact ratio from bulk dense phase CG simulation comparing conditions with and without RNA (ratio = +RNA/-RNA). Horizontal and vertical dashed line indicates the LC-RGG1 domain boundary. I) Representative simulation snapshot from bulk dense phase CG simulations of FUS LC-RGG1 and RNA. The LC domain is shown in blue, RGG1 domain in orange, and RNA in red. Bottom panel shows a magnified view of RNA-protein interactions, revealing that RGG1 domains are in close proximity to RNA, while LC domains exhibit enhanced homotypic interactions.

Towards our goal of studying the interactions between FUS LC-RGG1 and RNA in condensed phase, we next sought to quantify the extent of RNA incorporation in FUS condensed phases. We created condensates with a fixed protein concentration/amount mixed with increasing amounts of RNA and then analyzed RNA content by re-dissolving pelleted FUS – polyU RNA condensates in urea, measuring both protein and RNA concentrations using UV spectrophotometry. As expected, increasing the total RNA concentration resulted in greater incorporation of RNA into the condensed phase (Fig. 1D). However, the fraction of RNA partitioning into the condensed phase decreased with higher RNA levels. Importantly, direct concentration measurements via UV spectrophotometry of urea dissolved condensed phase samples showed lower protein (and total biomolecule) concentrations in RNA-containing condensates (Fig. 1E).

To further investigate the molecular basis of RNA partitioning into FUS LC-RGG1 condensates, we performed coarse-grained (CG) coexistence simulations with varying amounts of RNA mixed with FUS LC-RGG1 (Fig. 1F). Consistent with our experimental observations, high RNA concentrations destabilized the condensates, increasing the protein saturation concentration and reducing the dense phase concentration (Fig. 1F, Supp Fig. S1A). In agreement with experimental measurements, CG simulations showed that as total RNA increased, more RNA partitioned into the dense phase while protein concentration decreased (Fig. 1F). However, the negative slope of the tie line at elevated RNA levels indicated that this recruitment became less efficient. This trend reflects competing forces: while strong protein-RNA interactions could initially drive RNA incorporation, electrostatic repulsion between RNA chains eventually limits further partitioning. Upon further examination of the molecular interaction network within simulated bulk dense phase (see Methods), we observed that RGG1 domain formed comparatively stronger interactions with RNA compared to the LC domain (Fig. 1G). Additionally, as RGG1 became increasingly occupied by RNA at higher concentrations, LC domains compensated by forming more homotypic LC-LC contacts (Fig. 1H, I, Supp. Fig. S1B). This reorganization of the intracondensate interaction network, therefore, can disrupt protein-protein contacts that drive protein phase separation (1). Overall, reorganization of the protein interaction network explains the observed decrease in its dense phase concentration (Fig. 1E, I, Supp. Fig. S1A) and supports a model where sufficiently high RNA levels progressively displace protein-protein interactions, ultimately leading to condensate disassembly.

### Comparing the location and effects of RNA interactions with FUS in the dispersed and condensed phases

Previously, we showed that RNA does not readily interact with FUS LC alone and does not induce its phase separation (43). However, when FUS LC and RGG1 are linked, we hypothesized that FUS LC may transiently form contacts with RNA in the dispersed phase, the condensed phase, or both. To probe this, we studied the details of FUS-RNA interaction with residue-to-residue resolution in both the dispersed and condensed phases. We employed solution NMR under reentrant phase conditions with excess RNA. By comparing HSQC spectra of ^15^N labeled FUS with and without the addition of polyU RNA (Fig. 2A), we observed RNA-induced chemical shift perturbations (CSPs) primarily in the RGG1 domain, with some much smaller chemical shifts in the LC domain (Fig. 2B). We also compared these chemical shift perturbations with polyU to torula yeast RNA and observed similar CSP patterns which suggests non-specific RNA binding involves the same set of residues in the RGG1 domain regardless of RNA composition or sequence (Supp. Fig. S2A). Given that FUS LC-RGG1 forms substantial self-interaction even under non-phase-separating conditions in the absence of RNA (1), we hypothesize that the small LC domain CSPs we observed with RNA may not stem from direct LC-RNA interaction, but rather from RNA occupancy of the RGG1 domain that redistributes LC-LC and LC-RGG1 self-interactions. To test this, we repeated the RNA titration experiment using isolated FUS LC or FUS RGG1 (Supp. Fig.S2B). Upon adding RNA, the isolated LC domain showed no significant CSPs, while the isolated RGG1 domain exhibited CSPs similar to those observed in the LC-RGG1 construct. Therefore, our results suggest that in the reentrant phase, with the presence of excess RNA that prevents phase separation, FUS-RNA interaction is localized primarily to the RGG1 domain, while FUS self-association involving the LC domain is altered indirectly due to the RGG1-RNA interactions.

**Figure 2.**
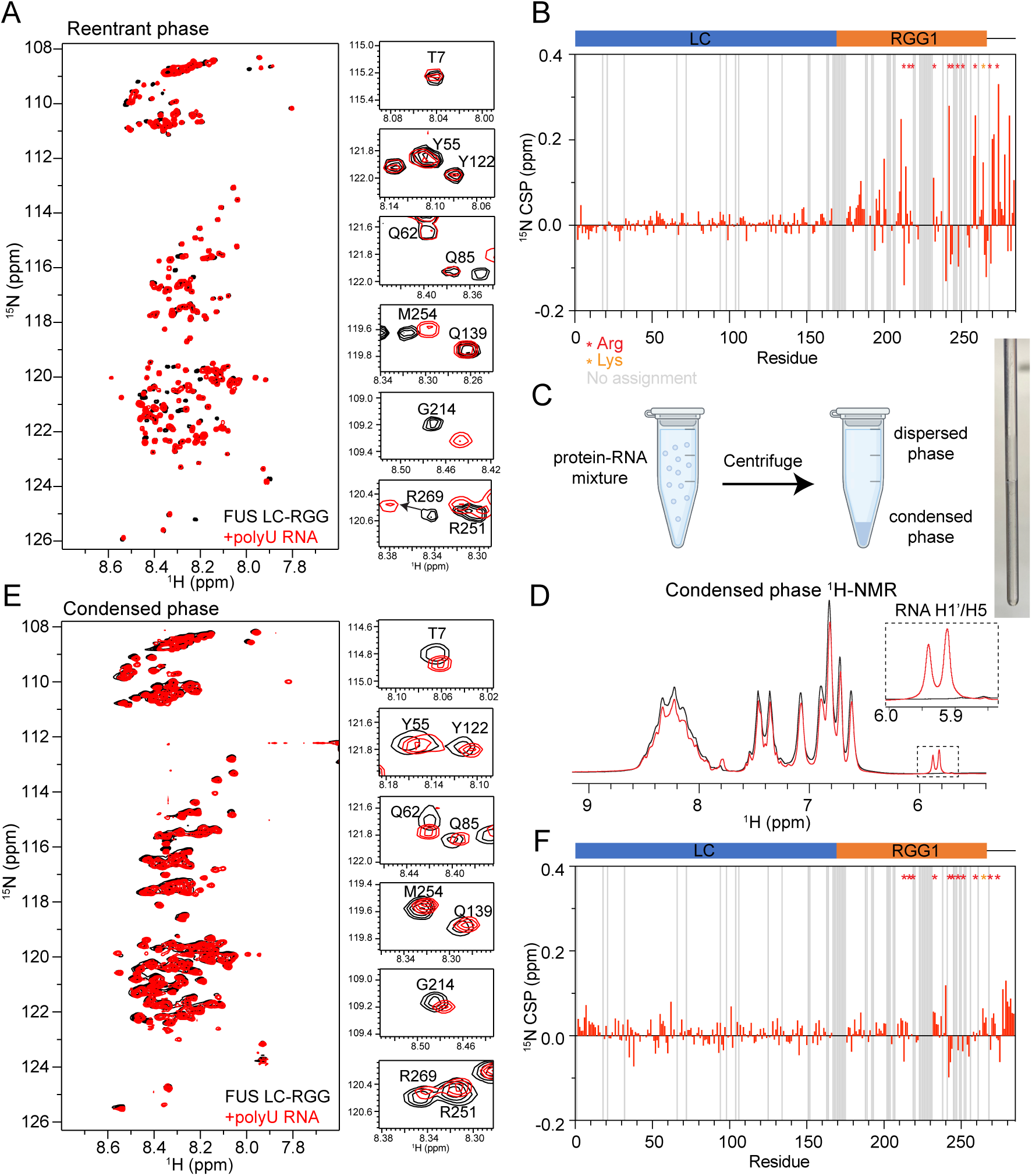
RNA induces distinct chemical shift perturbations in condensed-phase FUS. A) ^1^H-^15^N HSQC spectra of LC-RGG1 in the dispersed (no RNA) and reentrant (+polyU RNA) phases show chemical shift perturbations of residues primarily in the RGG1 domain. B) Quantification of ^15^N chemical shift perturbations from the spectra shown in A). C) Schematic of the workflow for preparing condensed-phase NMR samples. Phase separation was induced by mixing protein and RNA, droplets are consolidated by centrifugation, and the dense phase is carefully transferred into an NMR tube. D) 1D ^1^H spectra of FUS LC-RGG1 condensed phase after addition of polyU RNA (r*ed*) show RNA resonances not present without RNA (*black*). E) ^1^H-^15^N HSQC spectra of LC-RGG1 in the condensed phase, and examples of FUS ^1^H-^15^N HSQC peaks in absence and presence of RNA show chemical shift perturbations of residues across the entire protein. F) Quantification of ^15^N chemical shift perturbations in FUS LC-RGG induced by added RNA in the condensed phase.

We then sought to probe the molecular details of FUS-RNA interaction in the condensed phase and compare it to that in the dispersed phase. Specifically, we prepared pairs of macroscopic FUS condensed phase samples by inducing phase separation of FUS LC-RGG1 without and with polyU RNA and then centrifuging the resulting droplets to form macroscopic condensed phases (Fig. 2C). The obtained condensed phases, one containing approximately ≈10:1 (w/w) protein:RNA, were transferred to an NMR tube, as previously described (1). Although the ^1^H NMR signals of disordered proteins and RNA largely overlap, 1D ^1^H spectra collected on the FUS-RNA condensed phase revealed a set of additional resonances at 5.8-5.9 ppm corresponding to H1 and H5’ positions from RNA base and ribose (Fig. 2D), confirming the incorporation of RNA in the condensed phase.

We then compared 2D ^1^H-^15^N HSQC spectra of FUS LC-RGG1 condensed phase samples with and without RNA to probe the effect of the latter on the condensed phase (Fig. 2E). In contrast to the primarily RGG1-specific CSPs observed under re-entrant conditions (Fig. 2B), CSPs of comparable magnitude were observed in both the LC and the RGG1 domains in the condensed phase (Fig.2F). The maximum RNA-induced CSPs are smaller in the condensed phase compared to the dispersed phase (Supp. Fig. S2C), likely due to the much higher ratio of RNA to protein in the re-entrant dispersed phase. Interestingly, at individual residue level, several LC residues that did not shift upon RNA addition in the dispersed phase titration, such as T7, Y55, Q62, Q85, Q139, Y122, exhibited marked peak shifts in the condensed phase. Moreover, even for the RGG1 residues that displayed CSPs in both titrations, the direction of CSPs differed (e.g. G214, M254, R251, R269), suggesting distinct origins of the CSPs in the two phases (Fig. 2A, E). The CSPs between the dispersed phase without RNA and the re-entrant dispersed phase represent both the direct interaction of FUS with RNA (i.e. with RGG1) and changes to the FUS self-association as we described above. Despite a much lower protein to RNA ratio, the CSPs induced by addition of RNA in the condensed phase appear to arise from a reorganization of the protein interaction network and is likely associated with distinct changes to the condensate microenvironment and its internal dynamics, which we explore in the subsequent section.

### RNA modulates FUS LC-RGG1 dynamics in the condensed phase

To probe how RNA influence protein dynamics, we examined the effect of RNA on residue-specific motions using ^15^N relaxation NMR. In the dispersed phase, we observed that values of the transverse relaxation rate constant, *R*_2_ are elevated upon RNA addition for residues in the RGG1 but not LC (Fig. 3A), consistent with our observations that RNA interact primarily with RGG1 domain under these conditions. However, in the condensed phase we observed strikingly different behavior. Transverse (*R*_2_) relaxation rate constants (Fig. 3B), as well as longitudinal (*R*_1_) and the heteronuclear NOE (hetNOE) (Supp. Fig. S3A), showed widespread changes across the entire protein sequence in the RNA-containing condensate, in contrast to the dispersed phase where large changes were only seen in RGG1. The most prominent change after addition of RNA is decrease in *R*_2_ across most of the protein, consistent with overall faster molecular motions in the presence of RNA, an effect opposite to what is typically expected where stable binding slows reorientation.

**Figure 3.**
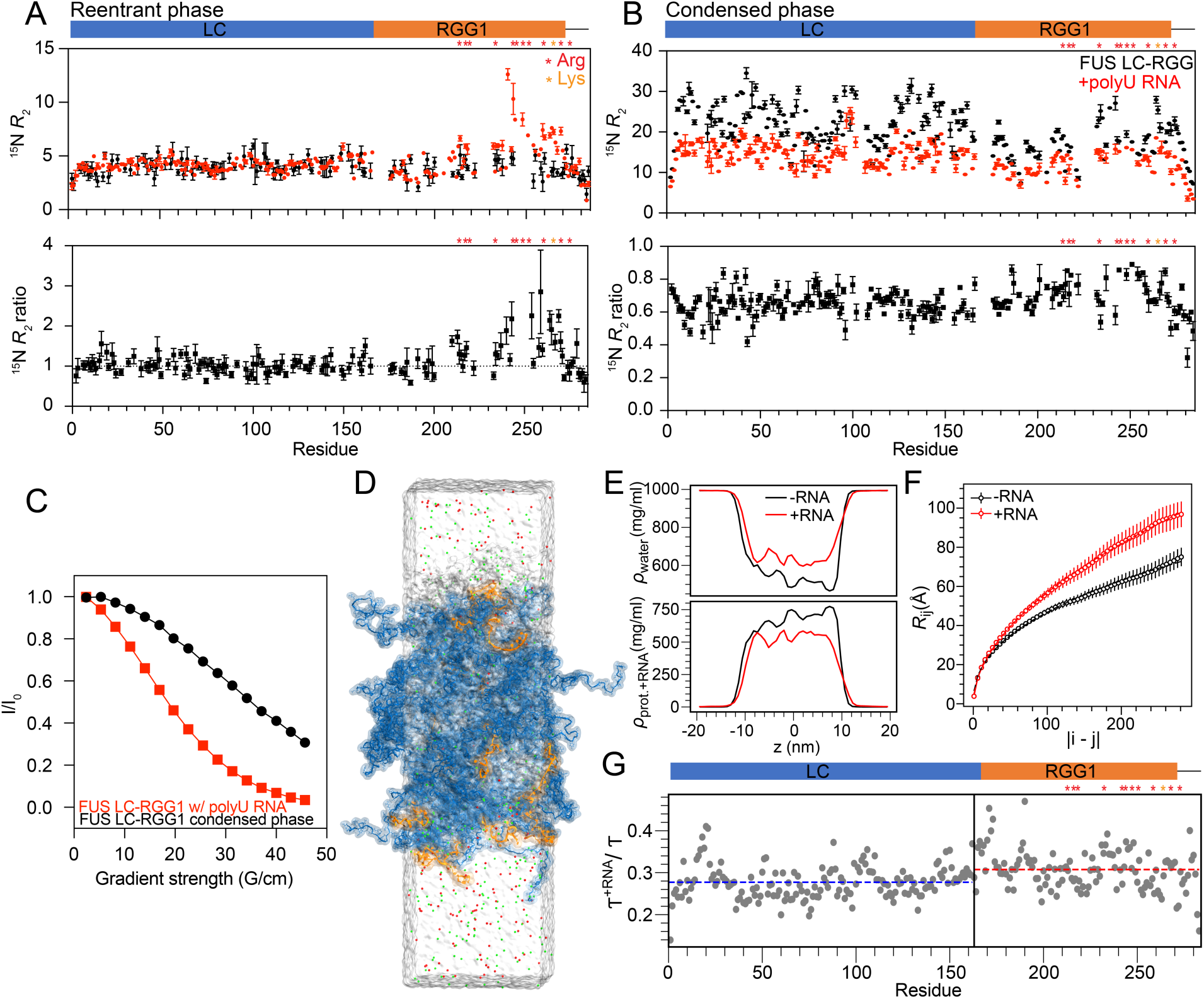
RNA accelerates FUS LC-RGG1 dynamics in the condensed phase. A) ^15^N *R*_2_ measurements of FUS LC-RGG1 in solution with and without added polyU RNA (top) and Δ*R*_2_ as result of the added RNA (*bottom*). These measures again show RNA-induced changes in the RGG1 region in solution. B) Condensed phase ^15^N *R*_2_ measurements with and without added polyU RNA (*top*) and Δ*R*_2_ as result of the added RNA (*bottom*) show changes consistent with faster motions with the largest changes found in the LC domain. C) Diffusion NMR measurements of FUS resonances in the condensed phases show faster diffusion of FUS in the presence of RNA. D) Snapshot from atomistic condensed phase simulation showing 25 FUS LC-RGG1 chains (blue), 18 U15 RNA chains (orange), Na+ (green) and Cl- (red) ions, and water (transparent surface). E) Concentration profiles of protein+RNA and water in presence and absence of RNA from condensed phase atomistic simulations. F) Inter-residue distances as a function of residue separation |i − j| for FUS LC-RGG1 in absence (black) and presence of RNA (red). G) Ratio of bond relaxation times for FUS LC-RGG1, calculated as the ratio of the relaxation time in the presence to that in the absence of RNA from condensed phase atomistic simulations. The horizontal dashed line in blue and red represents the mean value for LC and RGG1 domain respectively. Vertical line indicates the LC-RGG1 domain boundary.

We hypothesized that these relaxation changes are due to faster molecular motions arising from a less concentrated condensed phase when RNA is added (Fig. 1E). To test this, we measured the diffusion coefficient for the protein resonances within the condensates using pulsed field gradient NMR (Fig. 3C). Indeed, we observed that the protein diffusion in the RNA-containing condensate is 2.8x faster (D_app_ = 0.119 ± 0.009 μm^2^/s) than in its RNA-free counterpart (D_app_=0.042 ± 0.003 μm^2^/s), consistent with a more dynamic intra-condensate environment when RNA is added.

To investigate atomic-level details within condensed phase, we also performed atomistic simulations of FUS LC-RGG1 with RNA and compared the results to the RNA-free condensed phase (Fig. 3D). The simulations show decreased protein density and increased water content in RNA-containing phases (Fig. 3E), consistent with the experimental findings. Additionally, RNA incorporation into the condensed phase alters the ionic environment, increasing Na^+^ concentration while decreasing Cl^-^ concentration within the dense phase (Fig S3 B). The atomistic condensed phase simulations also suggest that, in the presence of RNA, the protein chains are more expanded, which could arise from heterotypic protein-RNA interactions (Fig. 3F).

Closer examination of the *R_2_* ratio profile in the condensed phase revealed additional insights (Fig. 3B). While *R_2_* values generally decreased in the presence of RNA—consistent with faster molecular motion—the reduction was slightly less pronounced in the RGG1 domain than in the LC domain. The bond relaxation times measured for atomistic condensed phase simulations also show that in the presence of RNA this orientational relaxation time decreases, indicating that the protein is more dynamic within the condensed phase, and that LC shows a lower relaxation time ratio value as compared to RGG1 (Fig 3G). We interpret this as a combined effect of increased global mobility due to phase dilution and persistent, domain-specific interactions between RGG1 and large, long chain RNA molecules. These direct RGG1 interactions with RNA partially counteract the overall decrease in *R_2_* in the RGG1 domain, which was also seen in the dispersed phase (Fig. 3A).

Together, these results demonstrate that RNA incorporation reduces condensate density and enhances overall protein mobility. However, the domain-specific relaxation patterns indicate that RGG1:RNA contacts still play an important and unique role in RNA binding within the condensed phase.

### Many types of amino acids mediate RNA contact in FUS-RNA condensed phase

Our results suggest that polyU RNA alters the molecular features of FUS LC-RGG1 condensates with domain/site specificity. To directly probe the interactions between FUS and RNA molecules within the condensed phase that give rise to these behaviors, we employed the ^13^C/^15^N double-half filtered ^1^H-^1^H NOE-HSQC experiments to detect intermolecular contacts between isotopically labeled protein and unlabeled RNA (44). In the ^13^C-edited NOE spectra, which resolve protein sidechain atoms involved in RNA binding, we observed robust NOE cross peaks linking RNA hydrogen atoms to a wide range of protein sidechain and backbone positions (Fig. 4A) including Arg HG, Ser HB, Gln HD, Tyr HD. Notably, the NOEs originate from RNA at positions H6, H1’/H5, and H3’/H4’, indicating contacts with both the nucleobases and the ribose backbone. Importantly, these cross peaks are absent in RNA-free control experiments, confirming their specificity for protein-RNA intermolecular interactions.

**Figure 4.**
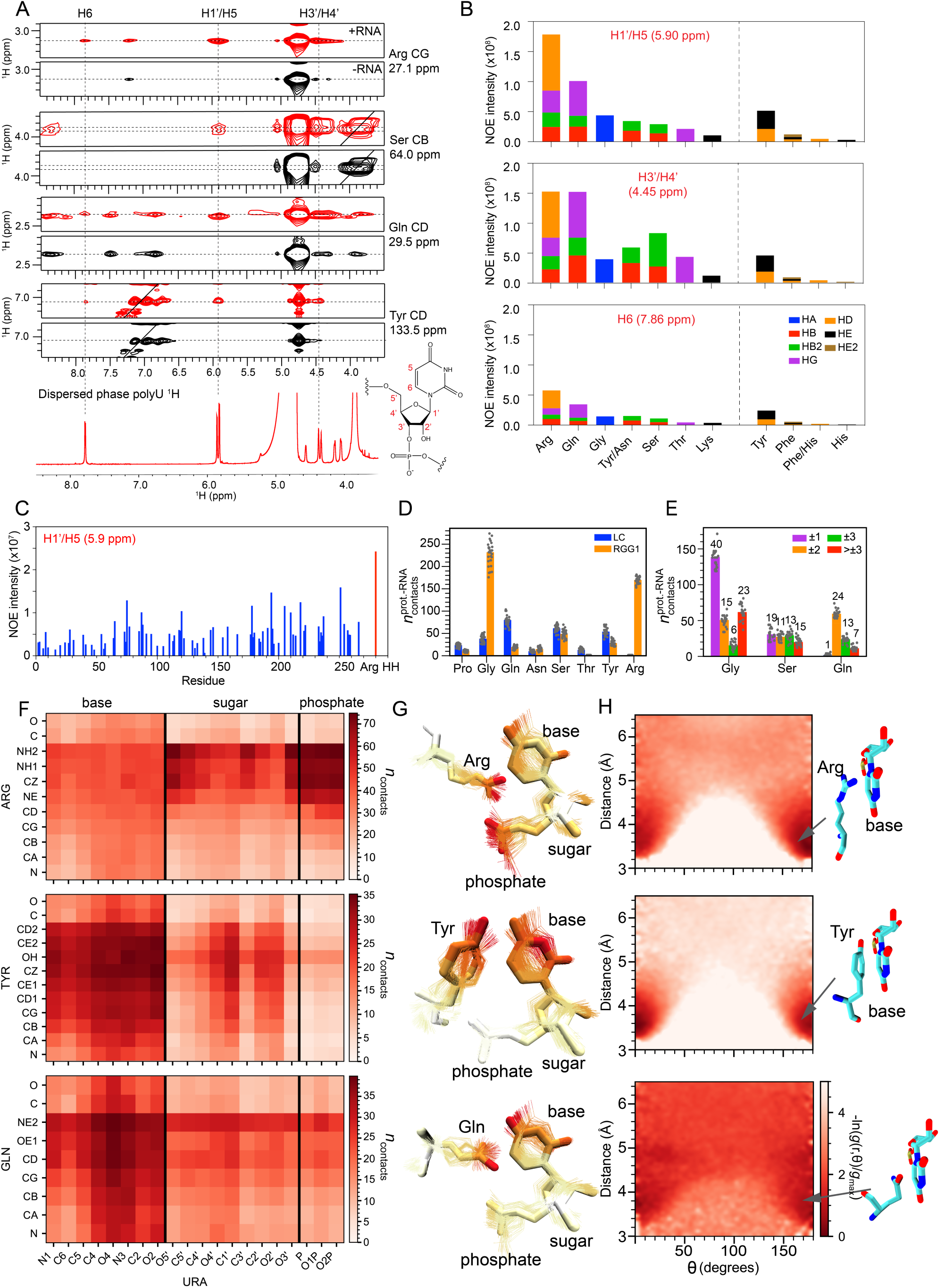
RNA forms interactions across FUS LC and RGG domains via many residue types. A) Intermolecular interaction between unlabeled RNA and ^13^C/^15^N double labeled FUS LC-RGG visualized using ^13^C/^15^N double-half filtered NOE-^13^C-HSQC. (left) A ^1^H-^1^H 2D projection of the experiment showed NOE from ^1^H resonances from RNA H2’/H3’/H4’/H5’ (4.45 ppm), H1’/H5 (5.90 ppm), and H6 (7.86 ppm) to protein sidechain protons, where the resonances present only in the +RNA condition (red) and not in the -RNA control (black) are true protein-RNA NOE contacts. (right) ^1^H-^13^C NOE plane for H1’/H5 at 5.90 ppm showed NOE to many types of amino acid sidechains. B) Quantifications of intermolecular NOE-^13^C-HSQC between RNA proton resonances and protein amino acid side chains. C) Quantifications of intermolecular NOE-^15^N-HSQC between RNA H1’/H5 and FUS LC-RGG amide backbone. D) Average RNA contacts per protein chain formed by abundant residue types from LC and RGG1 domains from atomistic condensed phase simulation. E) Average RNA contacts per protein chain formed by Gly, Ser, and Gln residues as a function of their sequence separation from the nearest Tyr or Arg (+/-1, +/-2, +/-3 positions, or >3 positions away) from atomistic condensed phase simulation. F) Contact map from atomistic condensed phase simulation showing heavy-atom proximities between FUS LC-RGG and RNA. G) Representative structures illustrating close contacts between RNA and selected amino acids (Arg, Tyr, Gln) from simulations. H) Contact orientation preferences between protein side chains and RNA bases in atomistic simulations of FUS:RNA mixtures, represented as potential of mean force (PMF) derived from pair correlation functions g(R,θ). Favorable stacking of aromatic RNA base and aromatic/planar amino acid side chains in of FUS:RNA mixtures appear as low PMF values near 0° and 180°. Corresponding molecular geometries are illustrated schematically on the right.

Quantitative analysis of NOE intensities at 5.9 ppm, which correspond to RNA ^1^H resonances at the H1′ and H5 positions, provided a mechanistic view of the FUS-RNA interaction (Fig. 4B). Consistent with previous studies highlighting the pivotal role of Arg residues in protein-RNA binding (45), Arg residues exhibited the strongest overall intermolecular NOE signals. Interestingly, we also detected significant NOEs for polar amino acid types not often associated with RNA interaction, including Gln, Gly, Tyr, Asn, Ser, Thr, and Phe (Fig. 4B). Since Tyr and Gln are primarily located in the LC domain, their involvement suggests extensive LC domain-RNA interaction within the condensed phase. Furthermore, NOE correlations at ^1^H positions specific to either sugar or base hydrogens revealed that the distinct chemical features of ribose and uracil moieties shape protein:RNA interaction patterns. At 4.45 ppm corresponding to the ribose H3’/H4’ positions, Gln and Ser sidechains exhibited more pronounced NOEs, likely due to interactions stabilized by hydrogen bonding with ribose hydroxyl groups. In contrast, at 7.86 ppm, corresponding to the H6 position of uracil bases, the pattern mirrored that of 5.9 ppm, with Arg showing stronger NOEs than all other residue types, emphasizing its dominant role in RNA base binding. Gly is represented in all of these positions, likely due to its prevalence in FUS sequence, accounting for nearly 30% of the FUS LC-RGG1, and are often found adjacent to other RNA-interacting amino acids. To further probe protein backbone-RNA interactions, we carried out the ^15^N-edited NOE experiments, which resolve NOEs to specific backbone positions along the protein (Fig. 4C). NOE signals at 5.9 ppm ^1^H position (RNA H1’/H5) were detected throughout the protein backbone, confirming extensive RNA contacts with both the LC and RGG1 domains. Notably, NOEs were slightly stronger for the RGG1 domain, consistent with interactions with the RGG1 in this phase, and in agreement with the large Δ*R*₂ in the RGG1 (Fig. 2E).

To provide additional details that are not visible by experiment, we investigated molecular level interactions in atomistic condensed phase simulations to characterize FUS LC-RGG-RNA interactions in the condensed phase. Importantly, the simulation observed the involvement of the same broad set of amino acid types observed experimentally, namely Gly, Arg, Ser, Gln, Tyr, in mediating RNA contacts (Fig. 4D, Supp. Fig. S4 A). While Gly initially appeared to be the dominant residue type interacting with RNA, normalization by residue abundance revealed that this dominance stems from Gly’s abundance in the sequence (Supp. Fig. S4B). Further analysis comparing RNA contacts of Gly and polar residues neighboring Tyr or Arg revealed that Gly’s apparent contact propensity stems primarily from its spatial proximity to these high-affinity RNA-binding residues (Fig. 4E).

The simulations also revealed that protein molecules interact with all major RNA components, base, sugar, and phosphate, with distinct binding preferences among amino acid types (Fig. 4F, G, Supp. Fig. S4 C,D). For example, Arg preferentially interacts with the phosphodiester backbone, consistent with predominant charge-charge interaction. Gln and Ser exhibit interactions through hydrogen bond-capable atoms, particularly Gln HE2/OE1 and Ser OG, with nucleotide carbonyl and hydroxyl groups, suggesting direct hydrogen bonding. In contrast, Tyr’s contacts are more evenly distributed across atoms within the planar uracil base. Though Gly’s contacts largely reflected its abundance and proximity to high-affinity residues, its backbone showed some propensity to interact with oxygen atoms of uracil bases (Supp. Fig. S4D).

To probe the spatial organization of these interactions, we analyzed the geometric orientations of protein-RNA contacts from atomistic simulations. Protein sidechain-uracil contact angle analysis showed probability basins near 0° and 180° at <4 Å (Fig. 4H), indicating preferred stacking geometries that were pronounced for Tyr and Arg (consistent with π–π and cation–π interactions, respectively), and weaker for Gln. Together, these findings indicate that diverse residues mediate condensed-phase protein-RNA contacts through π–π stacking, hydrogen bonding, and charge-charge interactions.

### Glutamine residues in FUS LC stabilize FUS-RNA co-phase separation

Our computational and experimental observations above strongly suggest that many residue types contribute to RNA interactions and specifically that interactions with the LC domain are important within the condensed phase. Although the requirement for positively charged residues like arginine and lysine for interaction with RNA has been established (14, 43, 46), we sought to verify the role of polar residue contacts to protein/RNA phase separation. Above, we noted that glutamine sidechain positions show strong NOEs, suggesting they are important contributors to FUS-RNA contacts. However, most glutamine residues are in the LC domain (37 out of total 45). To test the role of LC glutamine:RNA contacts, we used a glutamine-to-glycine (Q→G) variant of FUS LC-RGG, where all glutamine residues in the LC domain are replaced with glycine (Fig. 5A) (1). Despite extensive substitutions, we recently showed that the Q→G mutant retains nearly identical phase separation behavior as that of the wild-type (WT) protein (1), enabling us here to compare RNA partitioning at a similar extent of phase separation. We added polyU RNA to WT and Q→G FUS LC-RGG1 at 2:1 and 6:1 protein-to-RNA ratios under phase-separating conditions and analyzed the partitioning of both protein and RNA into the condensed phase. WT and Q→G FUS LC-RGG1 variants formed condensates to a similar extent at both conditions (Supp. Fig. S6). In contrast, a significant reduction in RNA incorporation was observed for the Q→G variant compared to the WT (Fig. 5B), which became more pronounced at higher protein concentrations. At a 6:1 protein-to-RNA ratio, approximately 64% of polyU RNA molecules partitioned into the WT FUS condensates, whereas only 43% the RNA entered the Q→G condensates. This result further supports a model where glutamine residues in the LC domain that lacks positively charged residues play a direct and critical role in recruiting RNA into the condensed phase.

**Figure 5.**
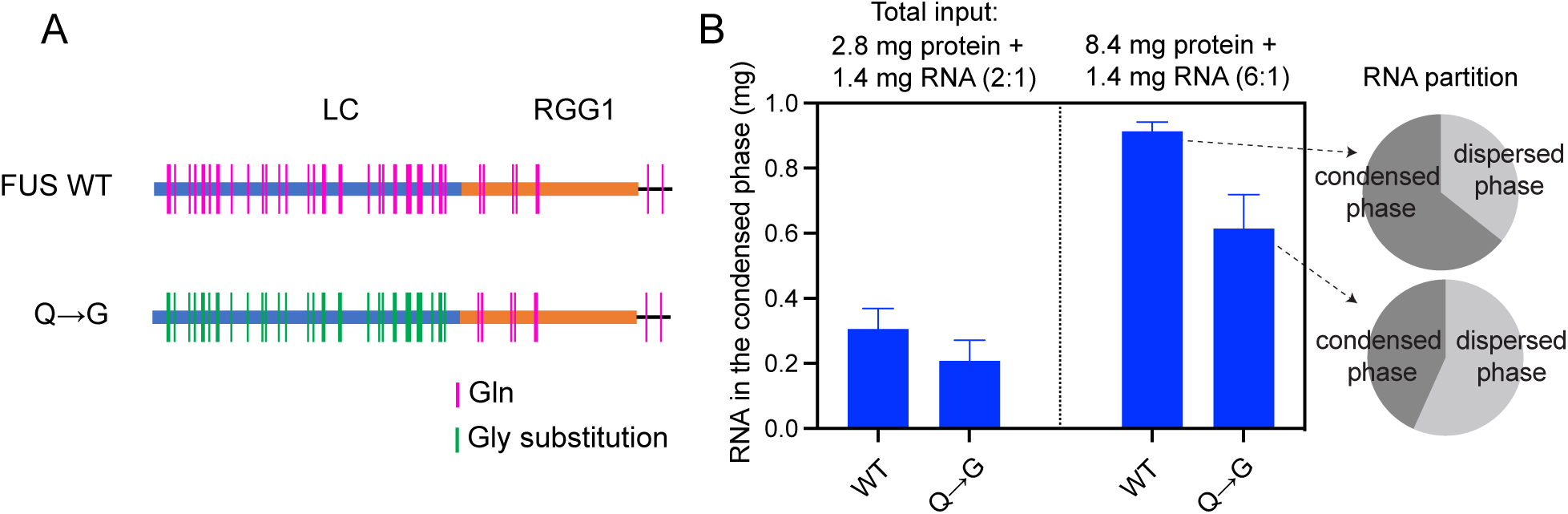
Glutamine residues stabilize FUS-RNA co-condensed phase. A) Glutamine residues are found in both the LC and RGG domains of FUS. The Q→G variant replaces all glutamine residues in the LC domain with glycine. B) RNA partitioning in the condensed phases formed by FUS WT or Q→G variant. (*left*) A fixed amount of 1.42 mg RNA was added to 2.8 mg or 8.4 mg FUS protein in 1 ml. The amount of RNA partitioned into the condensed phase was measured and plotted. (*right*) the fraction of total RNA found in the condensed and dispersed phase showed decreased RNA partitioning in the condensed with the mutant.

**Figure 6.**
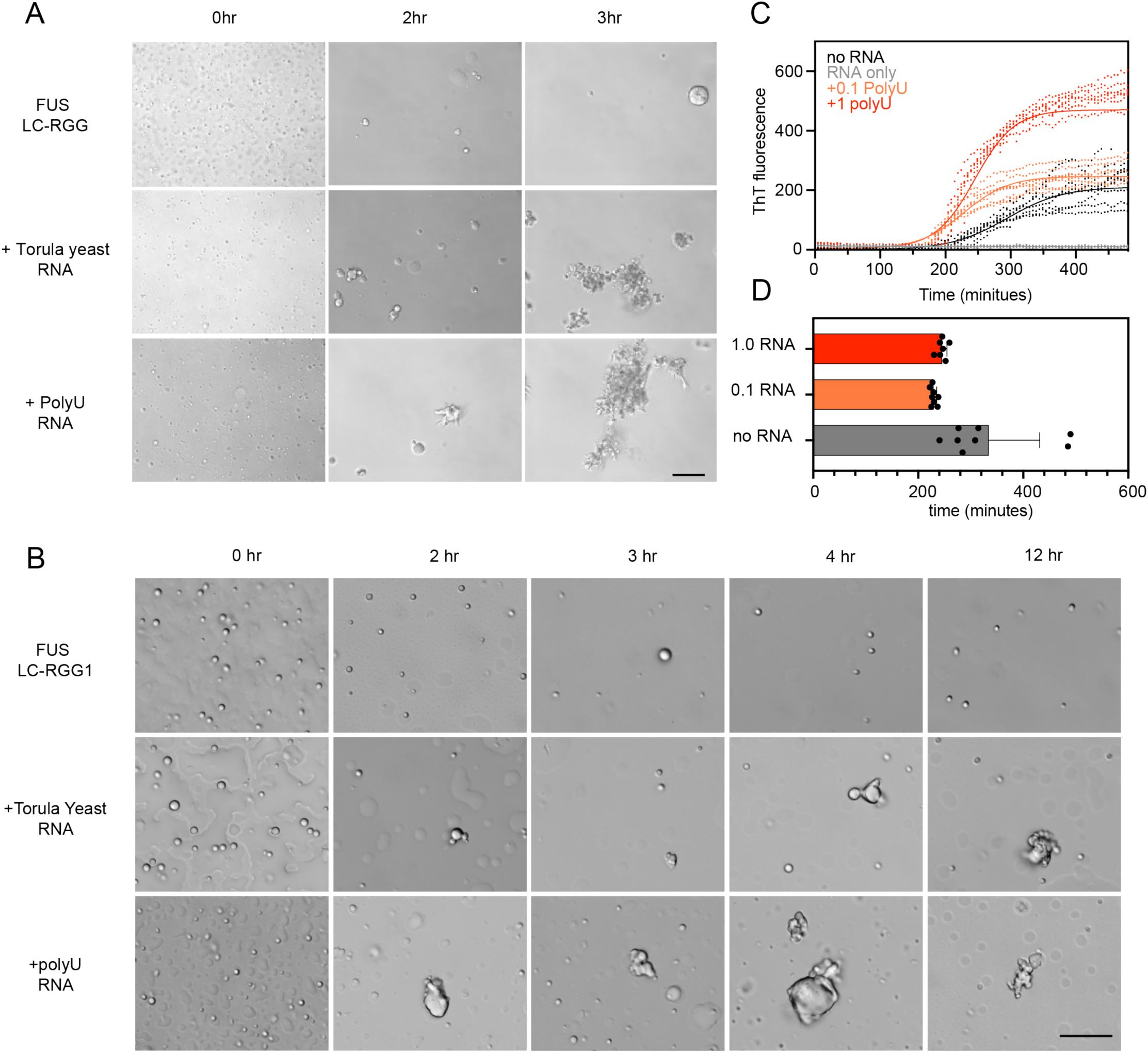
RNA accelerates FUS LC-RGG1 liquid-to-solid transition. B) DIC microscopy images of 50 μM FUS LC-RGG1 with and without adding 1:1 RNA (w/w, 1.4 mg/mL) (yeast or polyU RNA) after 0-3hr shaking agitation. C) DIC microscopy images of 50 μM FUS LC-RGG1 with and without adding 1:1 RNA (w/w) (yeast or polyU RNA) over 12 hours under quiescent condition. D) ThT intensities measured for 50 μM FUS LC-RGG1 mixture with polyU RNA at 1:0, 1:0.1 and 1:1 ratio as a function of time. The samples were subjected to shaking agitation in between readings. E) Time until ThT positive transition for the different samples in B).

### RNA accelerates FUS LC-RGG1 liquid-to-solid transition

Liquid-like droplets of full-length FUS condensates are well known to convert to solid-like assemblies consisting of β-sheet amyloid fibrils over time (47, 48), a process that is exacerbated by ALS/FTD-associated disease mutations in either LC or RGG domains that perturb RNA-binding (49). FUS aggregation is also influenced by RNA, while unstructured RNA is generally considered to be condensate-solubilizing, some reports have indicated that RNA sequences which form G-quadruplex structures can enhance the aggregation of full-length FUS (50). Other studies have suggested that RNA alters the properties of the surfaces of protein condensates, where protein aggregates may be nucleated (47, 51, 52).

Here we sought to investigate how RNA influences the liquid-to-solid transition of FUS condensates. Using DIC microscopy to monitor FUS LC-RGG1 condensates over time, we compared FUS LC-RGG1 samples with and without of RNA at a 1:1 (w/w) ratio under conditions where FUS LC-RGG1 is phase separated both in the absence and presence of RNA (50 μM) (Fig. 5A). Strikingly, addition of RNA, either yeast total extract RNA or polyU RNA, markedly accelerated the appearance of visible aggregates, both under shaking and quiescent conditions (Fig. 5A, B). RNA-containing samples transitioned from liquid-like droplets to increasingly solid aggregates far more rapidly than FUS-only controls.

To quantify the progression of the liquid-to-solid transition, we monitored thioflavin T (ThT) fluorescence, a sensitive readout for β-sheet-rich amyloid fibril formation (Fig. 5C). Both 1:0.1 and 1:1 FUS:RNA mixtures showed shorter delay times before ThT began to rise compared to FUS alone (Fig. 5D). Notably, this acceleration occurred not only at 1:0.1 protein-to-RNA, which promotes phase separation, but also at 1:1, where RNA shifts the system into the reentrant regime and reduces droplet formation. Moreover, the higher RNA condition reached substantially greater maximum ThT intensities. Taken together, these results indicate that RNA can enhance fibrillar aggregation of FUS LC-RGG1.

While the above findings may initially appear to be at odds with the known solubilizing effect of unstructured RNA, our results can be reconciled based on a model wherein RNA promotes a reorganization of the protein interaction network within the condensate to favor nucleation of amyloid aggregation. Specifically, our NMR and MD data indicate that RNA binding to RGG1 competes with RGG1-mediated contacts, thereby shifting the interaction balance toward increased LC–LC contacts (Fig. 1H), which ultimately promotes amyloid aggregation.

## Conclusion

Consistent with prior studies on RNA-binding proteins such as TDP-43 and hnRNPA1, our results demonstrate that RNA serves as a critical modulator of FUS phase behavior (36–39). We show that at low RNA concentrations, RNA promotes FUS phase separation, while at higher concentrations, it inhibits condensate formation. This reentrant behavior reflects competing processes: while at low RNA concentrations strong protein-RNA interactions drive RNA recruitment that enhances effective protein valency, at high RNA concentrations electrostatic repulsion between negatively charged RNA chains limits partitioning and destabilizes the condensed phase. Our CG simulations show that although RNA introduces additional protein-RNA contacts, it simultaneously disrupts critical protein-protein interactions, especially those between RGG-RGG and RGG-LC domains, which we have previously shown to be important for FUS phase separation (1). As RNA concentration increases, these protein-RNA interactions may become dominant, interfering with protein self-association and leading to disassembly of the condensates.

At the molecular level, our NMR data revealed that FUS-RNA interactions are predominantly mediated by the RGG1 domain in the dispersed phase, aligning with the established role of RGG motifs in RNA binding. However, in the condensed phase, both the LC and RGG1 domains engage in RNA interactions with RGG1 maintaining strong RNA contacts as evidenced by domain-specific relaxation dynamics. This expanded interaction mode may contribute to the distinct RNA interactome of FUS when phase separated in cells (35). Intriguingly, RNA also alters the FUS LC-RGG1 condensed phase as we also observed a decreased density of the condensed phase upon RNA addition. The substantially lower phase density with incorporation of only relatively small amounts of RNA by mass fraction may be used by cells to alter material properties of phases and also may represent ways to engineer condensates - for example, a shift toward a lower-density phase could speed reaction rates within biomolecular condensates.

Our combined NMR and atomistic simulation identified multiple amino acids, most notably arginine, tyrosine and glutamine, to mediate RNA contacts within the condensed phase, highlighting the complexity of protein-RNA interactions in these environments. Our results indicate that a diverse range of intermolecular forces, including π–π stacking, hydrogen bonding, and electrostatic interactions with RNA bases, ribose, and phosphate groups, all contribute to stabilizing FUS-RNA condensates. Importantly, we highlight the role of several polar amino acid types not primarily recognized for RNA binding, including Tyr, Gln, and Ser. We validated the role of glutamine in driving RNA partitioning into FUS condensed phases, a feature likely shared by other Q-rich RNA-binding proteins such as TDP-43 and TAF-15. Therefore, additional interactions between RNA and polar residues provide a more complex picture on how RNA is incorporated into condensates with proteins and points to additional means (e.g. glutamine vs glycine residue content) to achieve partitioning selectivity and hence functional specificity in cellular condensates.

A growing body of work shows that RNA is a potent modulator of RNA-binding protein aggregation, tuning low-complexity RBPs’ conversion into solid aggregates. For example, RNA can either promote condensate-mediated fibrillization of hnRNPA1, or at higher RNA:protein ratios, maintain proteins in a soluble state (53, 54). For TDP-43, specific binding to GU-rich RNA suppresses oligomerization and aggregation and loss or weakening of RNA binding promotes aberrant phase transitions and pathological assembly(55–57). Additional evidence suggests RNA to mitigate the liquid-to-solid transition of RNA binding proteins, thereby delaying aggregation (58, 59). Here, we observed decreased protein concentration and enhanced motions within RNA-containing condensates which may allow greater fluidity, potentially counteracting condensate rigidification that may lead to protein fibril formation within the bulk condensed phase (60, 61). However, when we make samples with droplets suspended in buffer, RNA accelerated the formation of FUS fibrillar aggregates. These seemingly paradoxical findings are likely due to RNA-induced remodeling of the FUS interaction network. Specifically, while RNA causes reduction in overall protein-protein contacts in the condensed phase, this decrease primarily reflects loss of RGG-mediated contacts, whereas LC-LC interactions are enhanced (Fig. 1H). Exchanging LC:RGG contacts for LC:LC contacts is consistent with a model in which RNA binding sequesters the RGG domain and frees the LC domain to engage in intermolecular contacts that promote aggregation. Because we find that macroscopic condensates, unlike suspended droplets of FUS with RNA, remain liquid-like over long times, this pro-aggregating effect of RNA appears to be stronger at the condensate interface with the dispersed phase. Indeed, several studies report that protein aggregation may be enhanced at the condensate interface (62–64). Other studies have suggested that RNA can alter the surface properties of protein condensates (47). If RNA preferentially partitions to the unique environment of the condensate surface, RNA-induced aggregation may therefore be further amplified there. Overall, these results highlight the nuanced and context-dependent nature of RNA’s effects on phase behavior and aggregation of RNA-binding proteins.

## Acknowledgements

This work was supported by NIGMS R01GM147677 (to N.L.F.), Human Frontier Science Program RGP0045/2018 (to N.L.F.), grant 1845734 from the National Science Foundation (to N.L.F.), NIGMS grant R35GM153388 (to J.M.). T.Z. was supported in part by a Pape Adams Postdoctoral Award from the Carney Institute for Brain Science at Brown University and a Milton Safenowitz Postdoctoral Fellowship (23-PDF-629) from the ALS Association. Computational resources were provided by the Texas A&M High Performance Research Computing (HPRC). The content is solely the responsibility of the authors and does not represent the official views of the funding agencies.

