## Supplemental Figures for "RNA modulates FUS condensate assembly, dynamics, and aggregation through diverse molecular contacts"

### Supporting Information

#### 1. Supplementary Figures

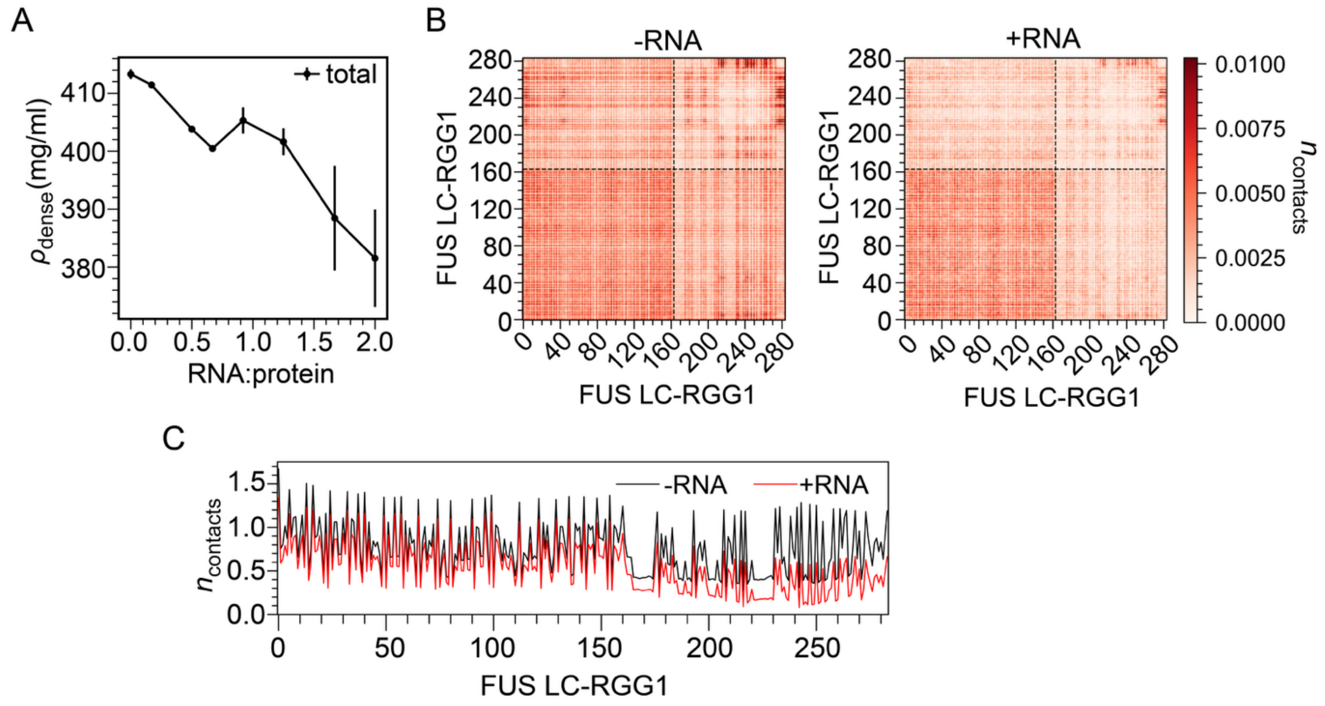


**Supp Figure S1. RNA Modulates Protein Interactions in CG simulation of FUS Condensates.**

A) Total concentration within dense phase as a function of increasing RNA concentration from CG co-existence phase simulations.

B) Per-chain intermolecular protein-protein contact map from CG bulk dense phase simulations in the absence (left) and in presence (right) of RNA.

C) Per-chain average number of intermolecular protein-protein contacts per residue from CG bulk dense phase simulations, in the absence and presence of RNA.

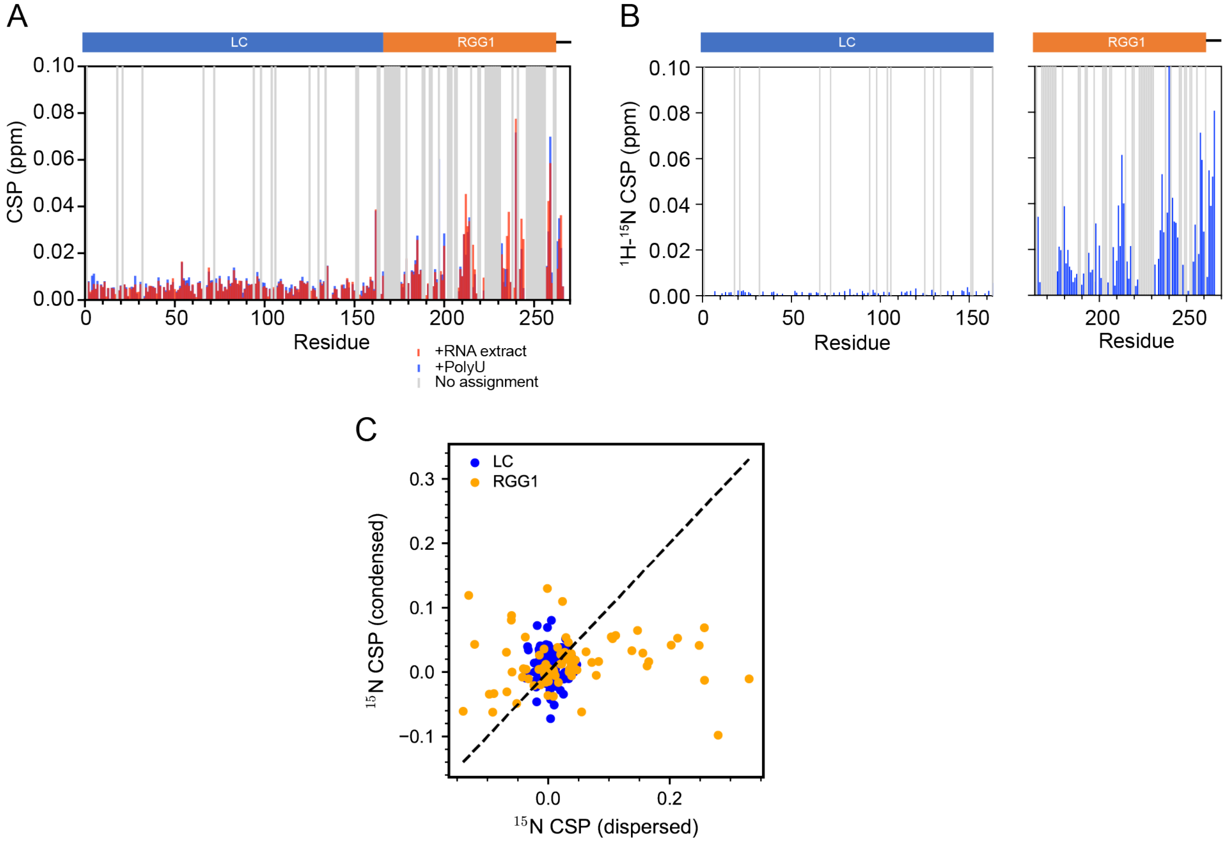


**Supp Figure S2. Isolated FUS RGG but not LC domain can interact with RNA.**

A) RNA induced $^1$H-$^{15}$N CSP for FUS LC-RGG1.

B) RNA induced $^1$H-$^{15}$N CSP for isolated LC or RGG1 domain.

C) CSP comparison for LC and RGG1 domains in dispersed and condensed phase.

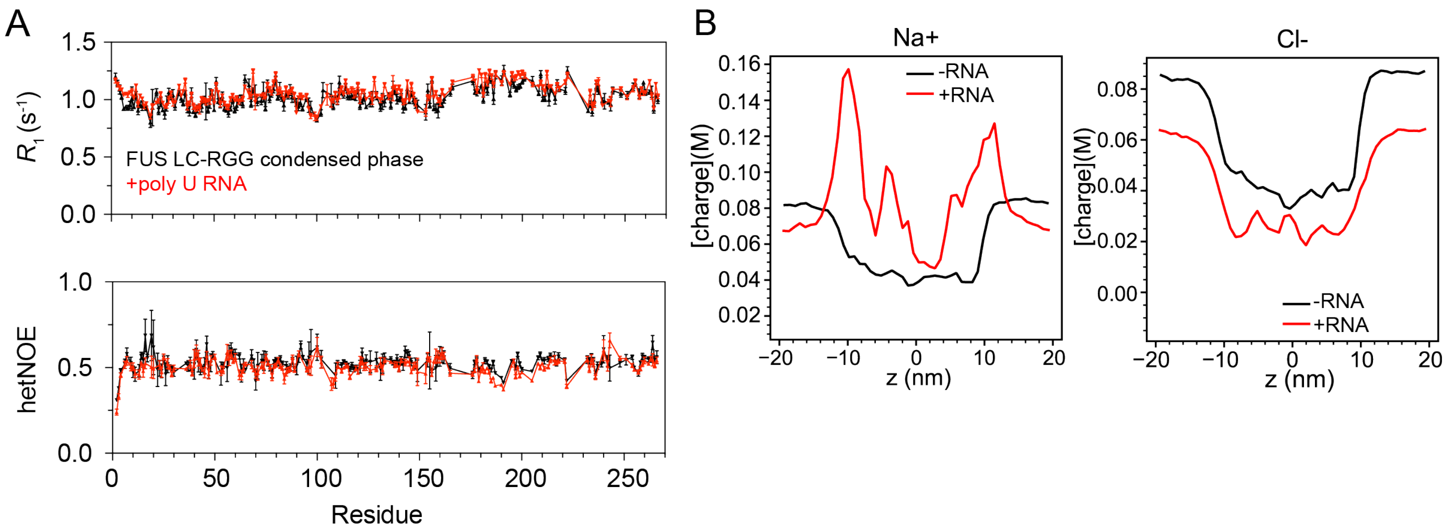


**Supp Figure S3. RNA incorporation changes FUS LC-RGG condensed phase**

A) $^{15}$N $R_1$ relaxation and hetNOE values for FUS LC-RGG1 in the condensed phase, in the absence and presence of polyU RNA.

B) Concentration profiles of ions in presence and absence of RNA from condensed phase atomistic simulations.

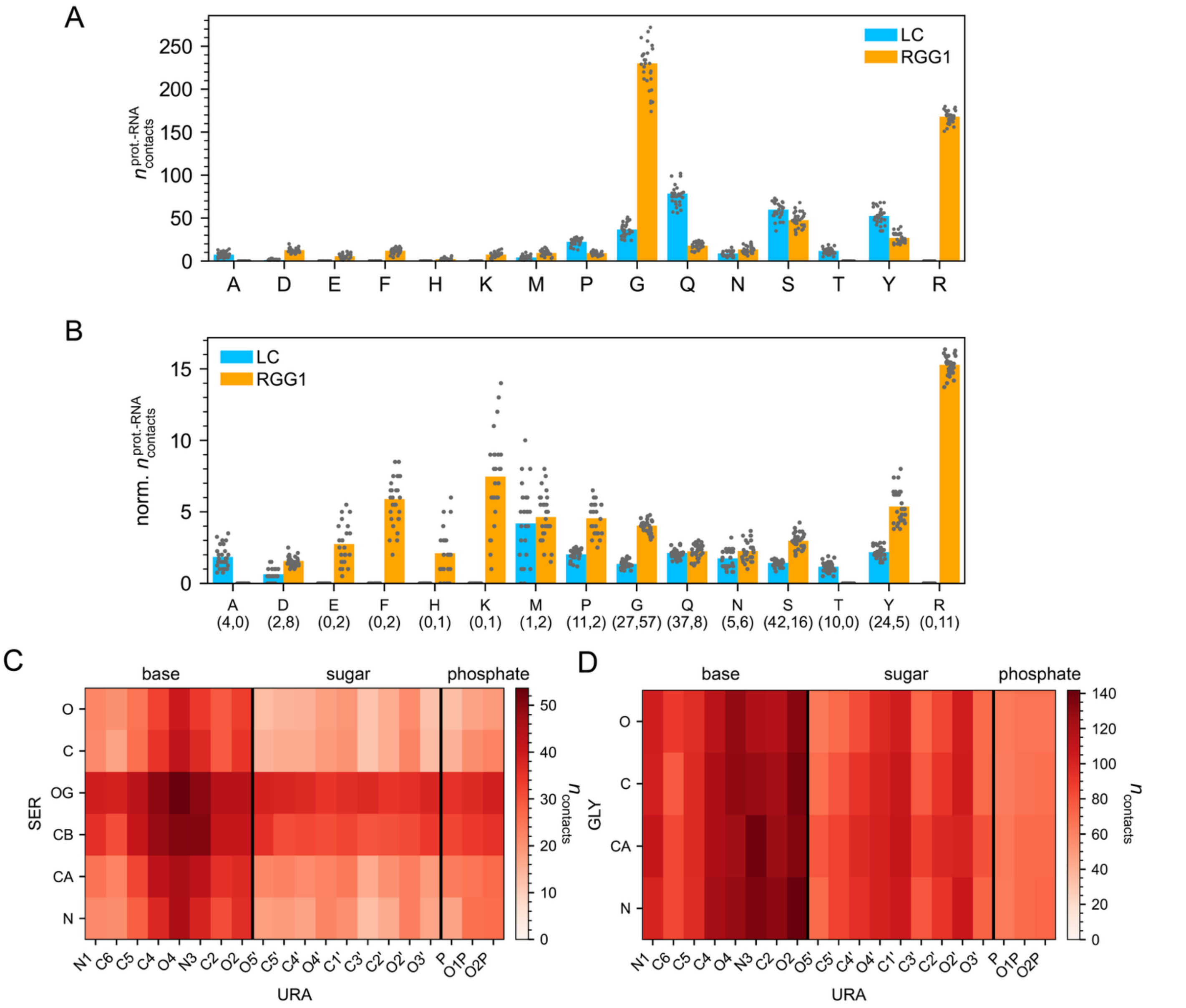


**Supp Figure S4. Many types of amino acids mediate RNA contact in FUS-RNA condensed phase**

A) Average RNA contacts per protein chain formed by all residue types from LC and RGG1 domains.

B) Average RNA contacts per protein chain formed by abundant residue types from LC and RGG1 domains normalized by their occurrence within each domain. The values below each residue type on the x-axis denote the occurrence of that residue type in the LC and RGG1 domains respectively.

C) Contact map showing heavy-atom proximities between Serine and RNA from atomistic condensed phase simulations.

D) Contact map showing heavy-atom proximities between Glycine and RNA from atomistic condensed phase simulations.

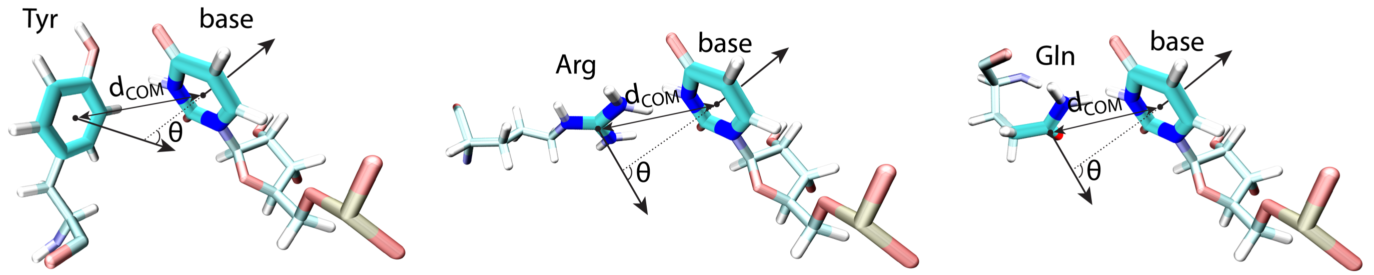


**Supp Figure S5. Schematic for contact angle computation.** Cartoons illustrate the planes of Tyr, Arg, and Gln used to compute the contact angle with the Ura base ring. The thick dark structures represent the selected planes. Angles were computed between the normals to the chosen planes, and the center-of-mass distances between the planes were used for contact angle analysis.

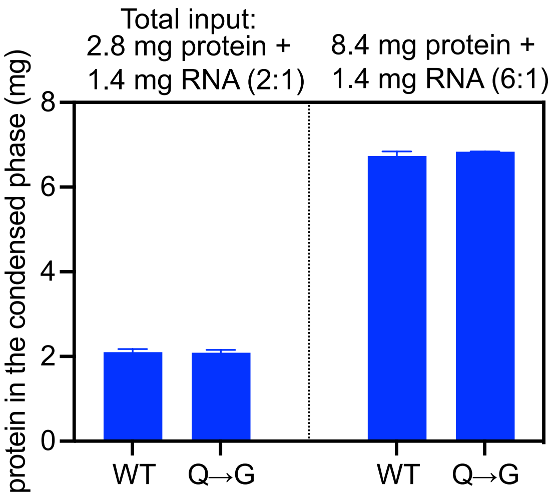


**Supp Figure S6. FUS LC-RGG1 Q→G variant phase separates similarly to WT**. A fixed amount of 1.42 mg RNA was added to 2.8 mg or 8.4 mg FUS protein in 1 ml. The amount of protein partitioned into the condensed phase was measured and plotted.

#### 2. Experimental procedures

##### Protein expression and purification

Hexahistidine-tagged human FUS LC-RGG1 (residues 1–284) was produced in *E. coli* BL21 Star (DE3) and purified from inclusion bodies as described previously (1), with the following procedure. Cells were cultured at 37 °C in M9 minimal medium containing 27 mM NaCl, 22 mM $KH_2PO_4$, 51 mM $Na_2HPO_4 \cdot 7H2O$, 1 mM $MgSO_4$, 1% v/v MEM vitamins, trace metals, 4 g/L glucose, and 1 g/L $NH_4Cl$. For isotope-enriched samples, $^{15}N$ $NH_4Cl$ was supplied as the sole nitrogen source. Protein expression was induced at an OD600 of approximately 0.8 with 1 mM IPTG (isopropyl β-D-1-thiogalactopyranoside) for 4 h at 37 °C. Cells were collected by centrifugation (7,800 g, 15 min, 4 °C), frozen at −80 °C, and subsequently resuspended in 20 mM sodium phosphate, 300 mM NaCl, pH 7.4 and lysed using a Fisherbrand Model 120 Sonic Dismembrator. The insoluble fraction was sedimented (47,855 g, 50 min, 4 °C), resuspended in the same buffer supplemented with 8 M urea and 10 mM imidazole, clarified by centrifugation again as above, and passed through a 0.2 µm filter. The solubilized fusion protein was captured by $Ni^{2+}$-affinity chromatography (5 mL HisTrap HP, Cytiva) under denaturing conditions with buffer A of 20 mM sodium phosphate, 300 mM NaCl, 8 M urea, pH 7.4 with 10 mM imidazole and eluted with the same buffer with 300 mM imidazole. Eluted fractions were combined and diluted ten-fold into 20 mM sodium phosphate, pH 7.4 to reduce the urea concentration prior to tag removal. $His_6$-TEV protease was added at a 1:50 to 1:100 w/w protease:substrate ratio, and the mixture was incubated overnight at room temperature. The reaction was then re-adjusted to 8 M urea and 10 mM imidazole and reapplied to the HisTrap column to remove the His-tag and $His_6$-TEV protease. The flow-through, containing tag-free FUS LC-RGG1, was buffer-exchanged into 20 mM CAPS, pH 11.0 to deprotonate tyrosine residues and minimize phase separation, concentrated using 3 kDa MWCO Amicon centrifugal filters, aliquoted, flash-frozen, and stored at −80 °C. Protein purity (>95%) was confirmed by SDS-PAGE.

##### NMR Sample preparation

All NMR buffers were 20 mM MES pH 5.5, 150 mM NaCl unless noted otherwise, with 5% $^2H_2O$ except for condensed phase samples with 10% $^2H_2O$ (to improve lock signal). Dispersed-phase FUS-RNA samples were made with $^{15}N$-labeled FUS LC-RGG1 at 25 µM, mixed with polyU at the concentrations described in the text. Samples were optically clear with no visible condensates. Condensed-phase samples were made as described earlier (1). Specifically, macroscopic condensates were generated by rapid ten-fold dilution of concentrated LC-RGG1 stock (~3 mM in 20 mM CAPS pH 11.0) into MES buffer at pH 5.5 with 150 mM NaCl and 10% $^2H_2O$, followed by centrifugation at 4,000 g for 10 min at 25 °C. For polyU RNA-containing condensates, RNA was added before centrifugation at a protein:RNA mass ratio of 10:1. After droplet formation, the condensed material was transferred into a 3 mm NMR tube and compacted by brief centrifugations at 1,000 g.

##### NMR spectroscopy

NMR spectra were acquired at 298 K on a Bruker Avance III HD 850 MHz spectrometer with a TCI HCN cryoprobe and z-gradients. Data were processed in NMRPipe and analyzed in CCPNMR 2.5.2. Water-suppressed $^1H$ spectra were collected with spectral width 16 ppm, TD = 32,768, and 16–128 scans. $^1H$-$^{15}N$ heteronuclear single quantum coherence (HSQC) spectra for the dispersed, biphasic and condensed phases were acquired with 3,072 direct time domain points and 400 indirect points with spectral widths spanning 10.5 ppm and 20 ppm in the direct and indirect dimensions, respectively. The $^1H$ and $^{15}N$ carrier frequencies were set to 4.7 ppm and 117 ppm, respectively, and direct and indirect acquisitions were collected for 172 and 116 ms each.

Triple resonance experiments, comprising HNCACB, CBCA(CO)NH, HNCO, HNCA, and HNN, were conducted to establish sequential and sidechain resonance assignments of FUS 1-284. Acquisition times for the triple resonance experiments were 185 ms in the direct $^1H$ dimensions, 24 ms in the indirect $^{15}N$ dimensions, 15 ms in the indirect CO dimensions, and 6 ms in the indirect Ca/Cb dimensions. Spectral widths were 13 ppm in $^{1}$H, 22 ppm in $^{15}$N, 7.5 ppm in CO dimension, and 54 ppm in Ca/Cb dimension.

Motions of the FUS LC-RGG1 backbone in the condensed and dispersed phases were measured using standard pulse sequences (hsqct1etf3gpsitc3d, hsqct2etf3gpsitc3d and hsqcnoef3gpsi). $^{15}$N longitudinal relaxation ($R_1$) was measured using a 7-point interleaved relaxation delay consisting of (100, 1,000, 200, 800, 300, 600 and 400 ms). Spectra at each delay value were acquired with 4,096 direct time domain points and 256 indirect points centered at 4.7 ppm and 117 ppm and with spectral widths of 10.5 ppm and 20 ppm for $^{1}$H and $^{15}$N, respectively. $^{15}$N transverse relaxation ($R_2$) was measured using a 7-point variable relaxation delay (16.9, 270.4, 185.9, 33.8, 118.3, 84.5 and 169 ms) using a 556 Hz CPMG field. Spectra at each point were acquired with 4,096 direct time domain points and 256 indirect points, carrier frequencies of 4.7 ppm and 117 ppm and spectral widths of 10.5 ppm and 20 ppm, respectively. Time domain data were acquired for 229 ms in the direct dimension and 74 ms in the indirect dimension. $^{15}$N heteronuclear NOEs were collected using interleaved steady-state NOE and no-NOE control experiments, each with a 5 s recycle delay. Spectra were collected with a center frequency of 4.7 ppm and 117 ppm, spectral widths of 10.5 ppm and 20 ppm and acquisition times of 172 ms and 149 ms for $^{1}$H and $^{15}$N, respectively.

Intermolecular NOE experiments (noesyhsqcgpwgx13d) to probe protein sidechain-RNA interactions were recorded on condensed phase samples containing 10:1 (w/w) mixture of $^{13}$C/$^{15}$N FUS LC-RGG1 with $^{12}$C/$^{14}$N polyU RNA. Aliphatic and aromatic regions of the spectrum were collected separately. For the aliphatic regions, a $^{13}$C carrier frequency of 43 ppm was used while the $^{1}$H carrier frequency was set to 4.7 ppm. Spectra were obtained with 4,096, 64 and 128 total points and spectral widths of 12, 80 and 12 ppm in F3, F2 and F1, respectively. A mixing time of 100 ms and 1 s recycle delay was used. For the aromatic regions, a $^{13}$C carrier frequency of 122.5 ppm was used with $^{1}$H carrier frequencies set to 4.7 ppm and the data were collected with the same spectral parameters as described above. Control samples with 100% $^{13}$C/$^{15}$N FUS LC-RGG1 and no RNA were created. Intermolecular NOE experiments (noesyhsqcf3gpwgx13d) to probe protein backbone–RNA interactions were recorded on the same condensed phase as above, containing 10:1 mixture of 13C/15N FUS LC–RGG1 and $^{12}$C/$^{14}$N polyU RNA. These data were collected with a $^{15}$N carrier frequency of 117 ppm and $^{1}$H carrier frequencies set to 4.7 ppm. Spectra were acquired with 4,096, 128 and 128 total points and spectral widths of 12, 20 and 12 ppm for F3, F2 and F1, respectively. A 100 ms mixing time and 0.8 s recycle delay were used. Control spectra were collected on 100% $^{13}$C/$^{15}$N FUS LC–RGG1 condensed phase samples. Data processing and analyses followed standard protocols using NMRPipe and CCPNMR Analysis 2.5 software (2, 3).

**Turbidity measurements**

Turbidity of LC-RGG1 with and without added RNA was monitored in 20 mM HEPES, pH 7.0, containing 150 mM NaCl at 25 °C. Samples were prepared by 10x dilution of FUS stocks in 20 mM CAPS, pH 11.0, directly into the assay buffer. Absorbance at 600 nm was recorded on a Cytation 5 plate reader. For each condition, individual samples were prepared, and 50 µl was transferred to a 96-well Costar plate and sealed with clear optical adhesive.

**ThT fluorescence assay**

To monitor aggregation of FUS LC-RGG1 samples, ThT fluorescence was recorded using a Cytation 5 plate reader as previously described (1). Eight replicate samples were prepared at 50 µM FUS LC-RGG1 by tenfold dilution of protein stocks in 20 mM CAPS, pH 11, into 20 mM MES, pH 5.5, 150 mM NaCl, and 20 µM ThT. Each sample (50 µl) was mixed thoroughly and transferred to a 96-well plate. Plates were incubated under double-orbital shaking at 807 cpm at 25 °C for 12 h. ThT fluorescence time courses were fit in GraphPad Prism 10 to a sigmoidal function of the form $y = d + a/(1 + \exp(-b(x - c)))$, where y is the ThT intensity, x is time, b is a parameter related to the apparent rate of ThT increase, c is the midpoint time of the transition, and d and a describe baseline and amplitude offsets. The inflection point c was taken as the characteristic aggregation time for each replicate.

**Microscopy**
DIC microscopy images of droplet and aggregates was obtained using a Nikon Ti2 microscope with 20x objective and 1.5x digital enhancement for all images. Phase separated samples were generated by 10x dilution from the CAPS storage buffer into 20 mM MES, pH 5.5, and 150 mM sodium chloride, mixed, and transferred to a glass cover slip for imaging. All images were processed using ImageJ.

**CG condensed phase simulations**
To understand how RNA modulates the phase behavior of FUS LC-RGG1, we performed CG coexistence phase simulations using CG models for both protein and RNA developed by our group (4). We simulated FUS LC-RGG1 using the HPS-Urry CG model while employing the HPS model for RNA (5). Similar to the protein CG model, the RNA CG model represents each nucleotide as a single bead. The diameter and hydropathy value of each nucleotide differentiate them in a manner analogous to how HPS-Urry parameterizes protein residues. The hydropathy values for each nucleotide were derived from partial charges on nucleotides in the OPLS all-atom RNA forcefield(6). Bonded and non-bonded interaction terms follow the same framework as for proteins, with the primary difference being an equilibrium bond length of 0.5 nm for RNA.
We simulated a fixed number of 60 FUS LC-RGG1 chains with varying numbers of polyU chains in a 15 × 15 × 105 nm slab geometry. Simulations were conducted with a 10 fs time step at 300 K. Temperature control was maintained using a Langevin thermostat with friction coefficient $\gamma = m_i/t_{damp}$, where mi represents residue mass and $t_{damp}$ was set to 1000 ps. The simulations were conducted using HOOMD-blue 2.9.3 (7). Each simulation ran for 5 µs, with the initial 1 µs excluded as equilibration time. Error bars were calculated by dividing trajectories into 4 blocks and computing the standard error of the mean.
The pairwise residue contact map was generated from bulk dense-phase simulations. These simulations were initially equilibrated in the NPT ensemble under an external isotropic pressure of P = 0 atm for 0.5 µs. Following the equilibration, 1 µs production runs were performed using Langevin dynamics at constant volume, keeping all other parameters the same as in the slab geometry simulations. Contacts were computed by considering two residues, *i* and *j*, to be in contact when their separation distance was less than 1.5σ. The contact propensity, $n_{contact}$, was determined as $\langle n_{ij} \rangle$, where $n_{ij}=1$ if the distance criterion was satisfied and $n_{ij}=0$ otherwise. In the resulting maps, contact propensities were expressed as $-\ln(n_{contact}/n_{max})$, where $n_{max}$ representing the highest contact propensity observed among all residue pairs.

**AA Condensed phase simulation**
For condensed phase simulations, we first performed CG simulations of protein and RNA to leverage the computational efficiency of CG models and obtain well-equilibrated structures. These were subsequently backmapped to all-atom configurations using established backmapping tools (8, 9). Initially, we simulated 25 FUS LC-RGG1 chains and 25 polyU chains (length = 15) in a box of 11 nm × 11 nm × 770 nm using the HPS Urry model for proteins and the HPS model for RNA.
After obtaining well-equilibrated structures, we selected protein and RNA within a defined Z-dimension (-25 nm to 25 nm), resulting in the selection of 25 protein chains and 18 RNA chains. We then extracted the protein configurations and backmapped them using MODELLER (9). RNA configurations were backmapped separately using Arena software (8). To enhance backmapping accuracy in the condensed phase, we employed a constrained protocol in MODELLER that better aligns all-atom structures with the original CG model. This protocol reconstructs side chains and backbone through two sequential steps. Initially, the CG model serves as a template to generate an initial configuration while optimizing only side chains. This approach ensures minimal displacement of $C_\alpha$ atoms, preserving the overall structure. Subsequently, the side-chain-optimized structure is used as a template for backbone refinement while keeping side chains fixed. This stepwise methodology prevents chain-level movement, maintains structural integrity, and significantly reduces steric clashes that occur during backmapping.

After obtaining atomistic structures of both protein and RNA, we concatenated the configurations and manually removed any clashes prepared the system in GROMACS v2021.3 (10) with box size of 11 nm × 11 nm × 40 nm. We used the Amber03ws (11) force field for proteins, which has been shown to capture protein folding thermodynamics and provides a more precise representation of the unfolded state. Moreover, it shows effectiveness in accurately representing both local and global conformational properties of IDPs and protein-protein interactions. For RNA, we used the χOL3 (12) force field with refined glycosidic torsion angles and adjusted phosphate oxygen radii (13). These modifications address known issues in RNA simulations, including the tendency to form unrealistic ladder-like structures and to suppress anti to high-anti χ shifts in RNA.

For both protein and RNA, interactions with water were optimized. For proteins, these optimized protein-water parameters have been shown to be useful for accurately capturing the conformational ensemble of IDPs (11). Tuning RNA–water interaction strengths has previously reduced artifactual base stacking in simulations of dye–ssRNA complexes (14). By applying the same approach to protein–RNA systems—i.e., strengthening RNA–water interactions—we anticipate suppressing unrealistically compact states characterized by excessive stacking and hydrogen bonding (15). This adjustment is intended to better capture the conformational dynamics of protein-RNA assemblies. However, a thorough, system-wide evaluation is still needed to validate this approach and quantify its effects on protein–RNA complexes under diverse conditions.

For system preparation, after placing protein and RNA chains in the box, we first performed vacuum minimization. The system was then solvated with TIP4P/2005 water (16), followed by energy minimization to relax any remaining steric clashes. The minimization steps were conducted using the steepest descent algorithm. Sodium and chloride ions were added to neutralize the charge and reach a salt concentration of 150 mM, using improved salt parameters proposed by Luo and Roux (17). Short temperature equilibration for 100 ps was carried out under the NVT ensemble at 300 K using a Nosé–Hoover (18) thermostat with a coupling constant of 1.0 ps. Short pressure equilibration to 1 bar for 100 ps was then performed using the Berendsen barostat (19) with isotropic coupling and a coupling constant of 5.0 ps.

Following system preparation in GROMACS, we conducted production runs in Amber22 by converting GROMACS topology and coordinate files to AMBER (20) input files (parm7 and rst7) using the "gromber" utility in ParmEd (21). In Amber, we implemented a multistep equilibration protocol. Energy minimization was performed using steepest descent for 10,000 steps followed by conjugate gradient for 20,000 steps. During this phase, protein and RNA chains were restrained with 100 kcal/mol/Å² on all heavy atoms. Following energy minimization, we heated the system from 100 K to 300 K over 2 ns with protein and RNA restrained at 100 kcal/mol/Å² on all heavy atoms. We then performed NPT equilibration with gradual restraint reduction on protein and RNA heavy atoms from 100 to 10 to 1 to 0.1 to no constraints. Each restraint level was simulated for 2 ns using a Monte Carlo barostat (22) with a 1.0 ps coupling constant and a Langevin thermostat (23) with a friction coefficient of 1.0 $ps^{-1}$. Anisotropic scaling was applied such that the box scaled in the z-dimension while x- and y-dimensions remained fixed. All equilibration steps up to this point used a 2 fs time step. After completing these steps, we performed a final 100 ns NPT equilibration at 4 fs time step. The SHAKE algorithm was employed to constrain hydrogen-containing bonds, long-range electrostatic interactions were computed using the PME method and van der Waals interactions were truncated at 0.9 nm (24–26). We conducted 2 μs simulations for the condensed phase. Through Rg autocorrelation analysis, we established a 300 ns equilibration period, which was excluded from subsequent analyses.

For contact analysis, a contact between residues *i* and *j* was defined based on non-bonded van der Waals interactions, calculated as the total number of heavy atom pairs (one from each residue) within 0.45 nm. This cutoff enables better representation of side-chain contributions, particularly for residues with longer side chains.

We quantified orientation preferences between protein side chains and the uracil base using methods described previously (27). Specifically, we constructed a joint pair-correlation function $g(R, \theta)$, defined over the centroid–centroid separation $(R)$ and the angle $(\theta)$ between the normals to the two reference planes. Reference planes were defined as the phenyl ring plane for Tyr, the guanidinium plane for Arg, the terminal amide plane for Gln, and the pyrimidine ring plane for uracil (Figure S5). From MD trajectory, all $(R,\theta)$ observations were accumulated into a uniformly binned 2D histogram and converted to $g(R,\theta)$ by applying the standard spherical-geometry normalization (dividing each bin by its shell volume proportional to $R^2 \sin\theta$), yielding a density per unit solid angle. Finally, we calculated the potential of mean force (PMF) as $-\ln(\frac{g(R,\theta)}{g_{max}})$, normalized such that the minimum value equals zero.
